# IMPPAT 2.0: an enhanced and expanded phytochemical atlas of Indian medicinal plants

**DOI:** 10.1101/2022.06.17.496609

**Authors:** R. P. Vivek-Ananth, Karthikeyan Mohanraj, Ajaya Kumar Sahoo, Areejit Samal

**Author notes:** Institute for Clinical Chemistry and Laboratory Medicine, Technische Universität Dresden, Dresden 01307, Germany (Karthikeyan Mohanraj). Corresponding author **Address for correspondence:** Areejit Samal, Computational Biology Group, The Institute of Mathematical Sciences (IMSc), CIT Campus, Taramani, Chennai 600113 India.

## Abstract

Compilation, curation, digitization and exploration of the phytochemical space of Indian medicinal plants can expedite ongoing efforts toward natural product and traditional knowledge based drug discovery. To this end, we present IMPPAT 2.0, an enhanced and expanded database, compiling manually curated information on 4010 Indian medicinal plants, 17967 phytochemicals, 1095 therapeutic uses and 1133 traditional Indian medicinal formulations. Notably, IMPPAT 2.0 compiles associations at the level of plant parts, and provides a FAIR compliant non-redundant *in silico* stereo-aware library of 17967 phytochemicals from Indian medicinal plants. The phytochemical library has been annotated with several useful properties to enable easier exploration of the chemical space. We also filtered a subset of 1335 drug-like phytochemicals of which majority have no similarity to existing approved drugs. Using cheminformatics, we have characterized the molecular complexity and molecular scaffold based structural diversity of the phytochemical space of Indian medicinal plants, and performed a comparative analysis with other chemical libraries. Altogether, IMPPAT is the largest phytochemical atlas of Indian medicinal plants which is accessible at: https://cb.imsc.res.in/imppat/.

## Introduction

Medicinal plants have been used for centuries to treat human ailments in different systems of traditional medicine across the world. Phytochemicals are the chemical factors behind the therapeutic action of such plants and the medicinal formulations prepared from them^1,2^. Consequently, significant research has been directed towards the identification of phytochemicals of medicinal plants^3–6^ to discover novel and biologically relevant molecules. Furthermore, phytochemicals along with other natural products represent a biologically relevant chemical space, produced by diverse organisms which have evolved to attain high level of fitness under varied selective pressures^7^. These aspects have rendered the natural product space as a key player in the identification and development of drugs against several diseases. This fact is cemented by the recent analysis by Newman *et al.*^8^ wherein the authors report that 34% of the small molecule approved drugs in the last four decades are either natural products, or natural product derived, or botanical drugs^8^. Still much of the natural product space remains largely unexplored providing significant scope for the identification of novel molecular scaffolds and fragments for the development of new drugs^8^.

Indian medicinal plants and their formulations have been used for ages in traditional Indian systems of medicine like Ayurveda and Siddha to treat a variety of human diseases^9^. These medicinal plants are a rich source of novel phytochemicals which can enrich and expand the natural product space. Much of the traditional knowledge on Indian medicinal plants, still largely remains buried in books and monographs. The non-digital nature of this information limits its complete and effective use in drug discovery research. Further, molecular mechanisms behind the therapeutic action of medicinal plants used in traditional Indian medicine remain largely undiscovered. This poses a significant challenge towards turning a largely experience-based enterprise to evidence-based practice, leading to modernization of traditional Indian medicine. In a nutshell, creation of a comprehensive database on Indian medicinal plants, their phytochemicals, their therapeutic uses and their traditional medicinal formulations will be of immense use in natural product and traditional knowledge based drug discovery.

Towards this goal, we had earlier built the manually curated database, IMPPAT (version 1.0)^10^, containing 1742 Indian <u>M</u>edicinal <u>P</u>lants, their 9596 <u>P</u>hytochemicals, <u>A</u>nd their <u>T</u>herapeutic uses. Importantly, IMPPAT 1.0 compiled two dimensional (2D) and three dimensional (3D) chemical structures of the 9596 phytochemicals in the database, along with their physicochemical, drug-likeness, and absorption, distribution, metabolism, excretion and toxicity (ADMET) properties. In short, IMPPAT 1.0 is the largest phytochemical atlas specific to Indian medicinal plants^10,11^ to date. Subsequent to publication, the IMPPAT phytochemical library has enabled several computer-aided drug discovery studies, including research on the identification of anti-SARS-CoV-2 drugs^12–17^.

Given the widespread use of IMPPAT 1.0, we have built IMPPAT 2.0, an enhanced and expanded phytochemical atlas of Indian medicinal plants (Figure 1). The latest update, IMPPAT 2.0, has built upon the published data of earlier version^10^, and now compiles information on 4010 Indian medicinal plants, 17967 phytochemicals, 1095 therapeutic uses and 1133 traditional Indian medicinal formulations (Table 1). We have highlighted the key features of IMPPAT 2.0 in Figure 1. Firstly, in IMPPAT 2.0, the coverage of the Indian medicinal plants is more than doubled, and the phytochemical and therapeutic use associations of the Indian medicinal plants have increased more than 5-fold in comparison with IMPPAT 1.0. Secondly, IMPPAT 2.0 now provides the phytochemical composition, therapeutic uses, and traditional medicinal formulations of Indian medicinal plants at the level of plant parts such as stem, root or leaves. Thirdly, through extensive manual curation and standardization, IMPPAT 2.0 provides a FAIR^18^ compliant non-redundant *in silico* stereo-aware library of 17967 phytochemicals with 2D and 3D chemical structures. Fourthly, we have characterized the molecular complexity and the molecular scaffold based structural diversity of the phytochemical space of IMPPAT 2.0, and thereafter, compared with other chemical libraries. Fifthly, we have also filtered a subset of 1335 drug-like phytochemicals using multiple drug-likeness rules. Finally, we have compared the phytochemicals in IMPPAT 2.0 with phytochemicals from Chinese medicinal plants. From our cheminformatics analysis, we find that phytochemicals in IMPPAT 2.0 are more likely enriched with specific protein binders rather than promiscuous binders, have scaffold diversity similar to many larger natural product libraries, and share minimum overlap with the phytochemical space of Chinese medicinal plants. These results highlight the uniqueness, utility and complementary nature of the phytochemical space of Indian medicinal plants captured in IMPPAT 2.0. IMPPAT 2.0 is accessible without any login or registration requirement via a user friendly web-interface at: https://cb.imsc.res.in/imppat/.

**Figure 1:**
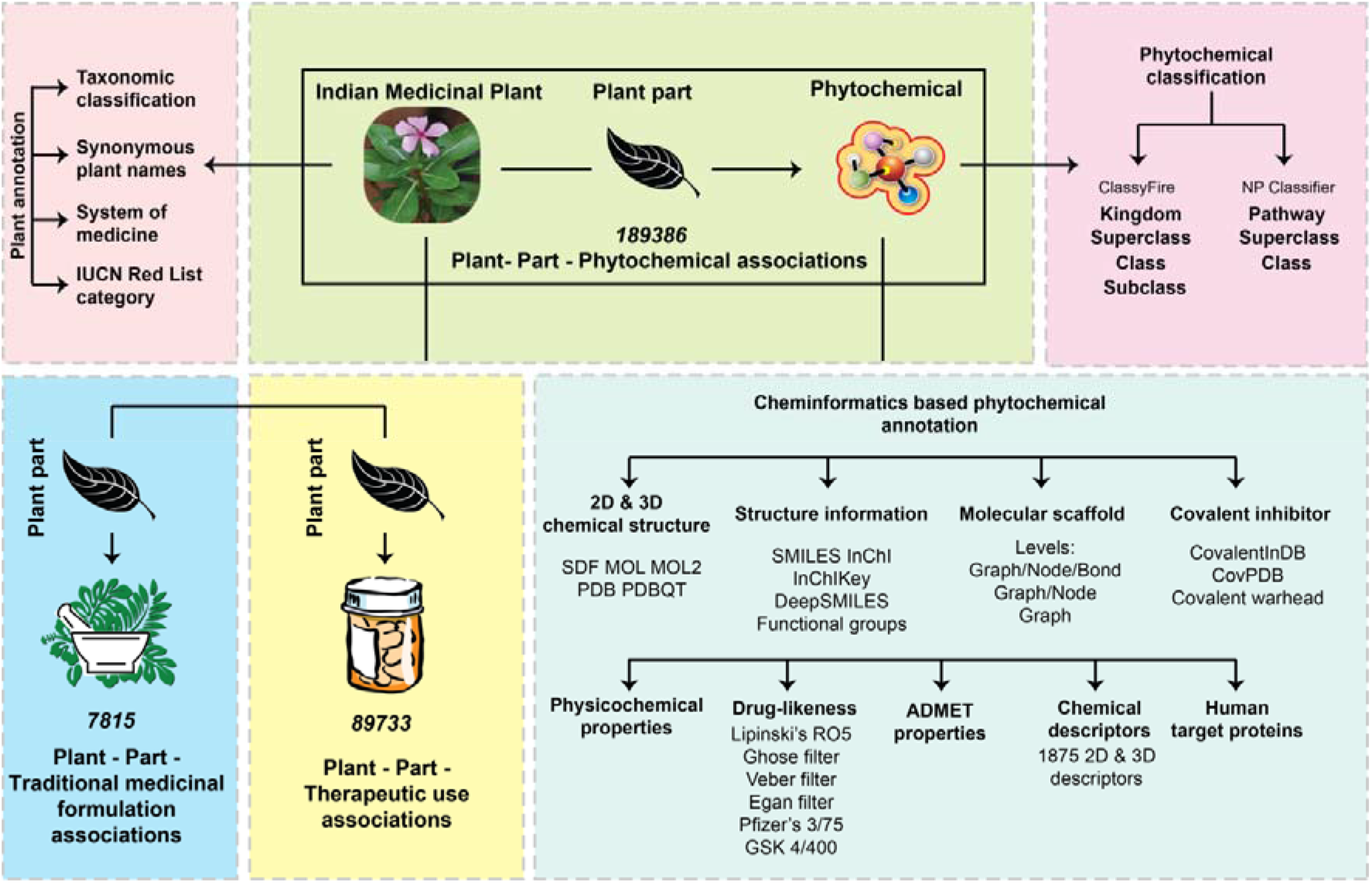
Schematic overview of the important features including enhancements and expansion realized in IMPPAT 2.0.

**Table 1:** Comparison of the updated version IMPPAT 2.0 with the previous version 1.0

| Feature | IMPPAT 2.0 | IMPPAT 1.0 |
| --- | --- | --- |
| Number of Indian medicinal plants | 4,010 | 1,742 |
| Number of Phytochemicals | 17,967 | 9,596 |
| Number of Plant - Part - Phytochemical associations | 189,386 | *Not available* |
| Number of Plant - Phytochemical associations | 124,995 | 27,074 |
| Number of Therapeutic uses | 1,095 | 1,124 |
| Number of Plant - Part - Therapeutic use associations | 89,733 | *Not available* |
| Number of Plant - Therapeutic use associations | 60,732 | 11,514 |
| Number of Traditional medicinal formulations | 1,133 | 974 |
| Number of Plant - Part – Traditional medicinal formulation associations | 7,815 | *Not available* |
| Number of Plant - Traditional medicinal formulation associations | 6,317 | 5,069 |

## Results

### Enhancement and expansion of IMPPAT

Previous version 1.0 of IMPPAT^10^ released in January 2018, is the largest online resource on phytochemicals of Indian medicinal plants. Here, we present the updated version 2.0 of IMPPAT, which is a significant enhancement and expansion over the previous version 1.0 (Table 1). This update was realized through extensive manual curation and addition of several new features to IMPPAT (Figure 1; Table 1). Figure 1 summarizes the important features including enhancements accomplished in IMPPAT 2.0.

### Increase in coverage of Indian medicinal plants

IMPPAT 2.0 compiles curated information on phytochemicals and therapeutic uses of 4010 Indian medicinal plants. The updated database achieves more than 2-fold increase in the coverage of medicinal plants with respect to the previous version (Table 1). During data collection from various sources, we encountered extensive use of synonymous plant names in published literature reporting information on phytochemicals and therapeutic uses of medicinal plants. This use of synonymous plant names can create difficulties while choosing the correct plant for phytochemical extraction or preparation of pharmaceutical formulations as prescribed in traditional medicine pharmacopoeia. For this reason, IMPPAT 2.0 provides the compiled information for a non-redundant list of 4010 Indian medicinal plants. This non-redundant list was created via an extensive manual curation effort as follows. First, we compiled a list of more than 7000 synonymous names corresponding to Indian medicinal plants for which the phytochemical information was collected from published literature in IMPPAT 1.0 or during this update. Second, The Plant List database (http://www.theplantlist.org/) was used to identify the accepted scientific names for the compiled plant names. Third, the synonymous names were merged using the accepted scientific names.

Further, the Indian medicinal plants covered in IMPPAT 2.0 have been annotated with information on their taxonomic classification, their use in traditional Indian systems of medicine, their synonymous names, and their present category in the IUCN Red list of threatened species^19^ (Methods). The 4010 Indian medicinal plants in IMPPAT 2.0 belong to 244 taxonomic families, and Figure 2a shows the families with more than 50 Indian medicinal plants in our database. In particular, Leguminosae is the largest family with more than 350 plants in IMPPAT 2.0. This is expected as Leguminosae, commonly known as legume, pea or bean family, is a large and medicinally important family of flowering plants^20^. The next two large families in IMPPAT 2.0 are Compositae and Lamiaceae, both of which are again families of flowering plants. Flowering plants or Angiosperms constitute 96% of the plants in IMPPAT 2.0. The remaining plants are Gymnosperms (2%) which include conifers and cycads, and Pteridophytes (2%) which include ferns and fern allies (Figure 2b). The medicinal plants captured in IMPPAT 2.0 are used in one or more traditional Indian systems of medicine such as Ayurveda, Siddha, Unani, Sowa-Rigpa and Homeopathy. In particular, 1328 plants in IMPPAT 2.0 are used in Ayurveda, followed by 1151 plants used in Siddha (Figure 2c). Precariously, we find many of the Indian medicinal plants require extensive conservation effort as 72, 50, 40, 11 and 3 plants are categorized in the IUCN Red list of threatened species^19^ as vulnerable (VU), near threatened (NT), endangered (EN), critically endangered (CR), and extinct in the wild (EW), respectively (Figure 2d).

**Figure 2:**
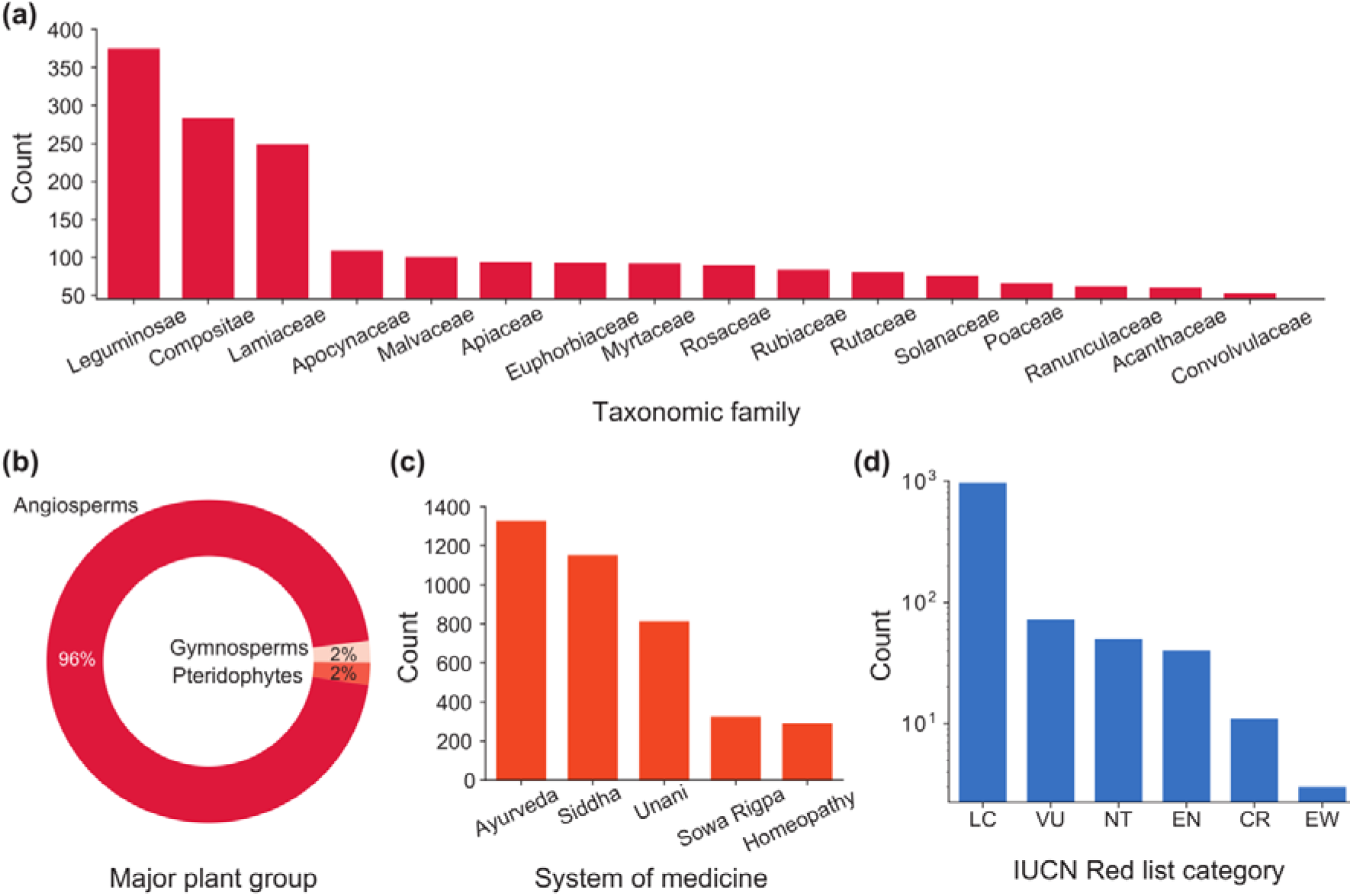
Coverage of Indian medicinal plants in IMPPAT 2.0. **(a)** Top taxonomic families of Indian medicinal plants in IMPPAT 2.0. Note only families with more than 50 Indian medicinal plants in the database are shown. **(b)** Classification of the Indian medicinal plants in IMPPAT 2.0 into major plant groups: Angiosperms (Flowering plants), Gymnosperms (Conifers, cycads and allies) and Pteridophytes (Ferns and fern allies). **(c)** Use of Indian medicinal plants in IMPPAT 2.0 in traditional Indian systems of medicine such as Ayurveda, Siddha, Unani, Sowa-Rigpa and Homeopathy. Note that a given Indian medicinal plant can be used in multiple systems of medicine. **(d)** Present category according to conservation status of the Indian medicinal plants in IMPPAT 2.0. LC – Least concern, VU – Vulnerable, NT – Near threatened, EN – Endangered, CR – Critically endangered, EW – Extinct in the wild.

### Information at the level of plant parts

Unlike previous version 1.0, IMPPAT 2.0 provides information on plant – phytochemical, plant – therapeutic use, and plant – traditional medicinal formulation associations at the level of plant parts (Table 1). For instance, the updated database compiles published information on the phytochemical composition for any Indian medicinal plant at the level of plant parts such as stem, root or leaves. Since it is common knowledge that phytochemical composition can significantly vary across different plant parts, this enhancement in IMPPAT 2.0 will facilitate researchers in phytochemistry and pharmacognosy to choose the appropriate protocol for extraction of the phytochemical of their interest for drug discovery studies. Moreover, traditional Indian systems of medicine, such as Ayurveda and Siddha, use specific plant parts for preparation of medicinal formulations used to treat various diseases. This further underscores the importance of compiled information in IMPPAT 2.0 on therapeutic use and traditional medicinal formulation at the level of plant parts.

### Increase in coverage of phytochemicals

Among the major enhancements in IMPPAT 2.0 is the creation of a non-redundant stereo-aware natural product library of 17967 phytochemicals specific to Indian medicinal plants. This represents a nearly 2-fold expansion in the size of the phytochemical library in comparison to the previous version 1.0 (Table 1).

Building upon the published methodology and extensive data compiled in IMPPAT 1.0^10^, we expanded the phytochemical associations in IMPPAT 2.0 as follows. First, the bulk of the plant – part – phytochemical associations for Indian medicinal plants were manually collected, curated and digitized from 70 specialized books (Supplementary Table S1). Only 9 out of these 70 books were covered in IMPPAT 1.0. Importantly, the remaining 61 books covered in IMPPAT 2.0 include: (a) 5 volumes of The Wealth of India published by the Council of Scientific and Industrial Research, Government of India, (b) 14 volumes of Ayurvedic, Siddha and Unani pharmacopoeias of India published by the Ministry of AYUSH, Government of India, and (c) 18 volumes of the Reviews of Indian medicinal plants published by the Indian Council of Medical Research (ICMR), Government of India. These valuable yet non-digitized book sources on Indian medicinal plants are known for their comprehensiveness and accuracy^21^. Second, aside from the books, all the plant – phytochemical associations compiled from various sources in the previous version IMPPAT^10^ 1.0 were manually revisited to additionally gather and curate phytochemical information at the level of plant parts. This last step also involved manual curation of more than 7000 research articles covered in IMPPAT 1.0 to gather additional information at the level of plant parts. Third, we incorporated data from a published database^22^ providing phytochemical information for the Indian medicinal plant *Rauvolfia serpentina*.

A major challenge during compilation, curation and digitization of the phytochemical composition of Indian medicinal plants is the large-scale use of non-standard and synonymous names for phytochemicals in books and research articles. Therefore, to create a non-redundant list of phytochemicals, we have standardized the phytochemical names fetched from diverse sources as follows. First, we mapped the chemical names to identifiers in standard databases such as PubChem^23^ and retrieved the associated two-dimensional (2D) and three-dimensional (3D) structures. Second, we compared the phytochemicals based on their structural similarity. Third, we manually checked the stereochemistry of the phytochemicals using the InChI. These steps led to the creation of a non-redundant stereo-aware chemical library of 17967 phytochemicals which are produced by 4010 Indian medicinal plants with therapeutic uses. Thus, the phytochemical atlas will aid ongoing efforts towards the identification of novel bioactive and therapeutic molecules.

Overall, there are 189386 non-redundant plant – part – phytochemical associations in IMPPAT 2.0 spanning 4010 Indian medicinal plants and 17967 phytochemicals. At the level of plant – phytochemical associations (after ignoring the plant parts), there is a 5-fold increase in IMPPAT 2.0 (Table 1). Figure 3a shows the occurrence of phytochemicals across 4010 Indian medicinal plants in IMPPAT 2.0. It can be seen that a majority of the phytochemicals (15335) have been reported to be produced by < 5 Indian medicinal plants, while a minority of the phytochemicals (114) are produced by > 200 Indian medicinal plants. In IMPPAT 2.0, *Psidium guajava* (468)*, Citrus sinensis* (457)*, Catharanthus roseus* (427)*, Coriandrum sativum* (403)*, Artemisia annua* (393)*, Rosmarinus officinalis* (391)*, Daucus carota* (391)*, Origanum vulgare* (366)*, Citrus reticulata* (364) and *Salvia officinalis* (363) are the top ten plants in terms of the compiled information on the number of phytochemicals produced by them.

**Figure 3:**
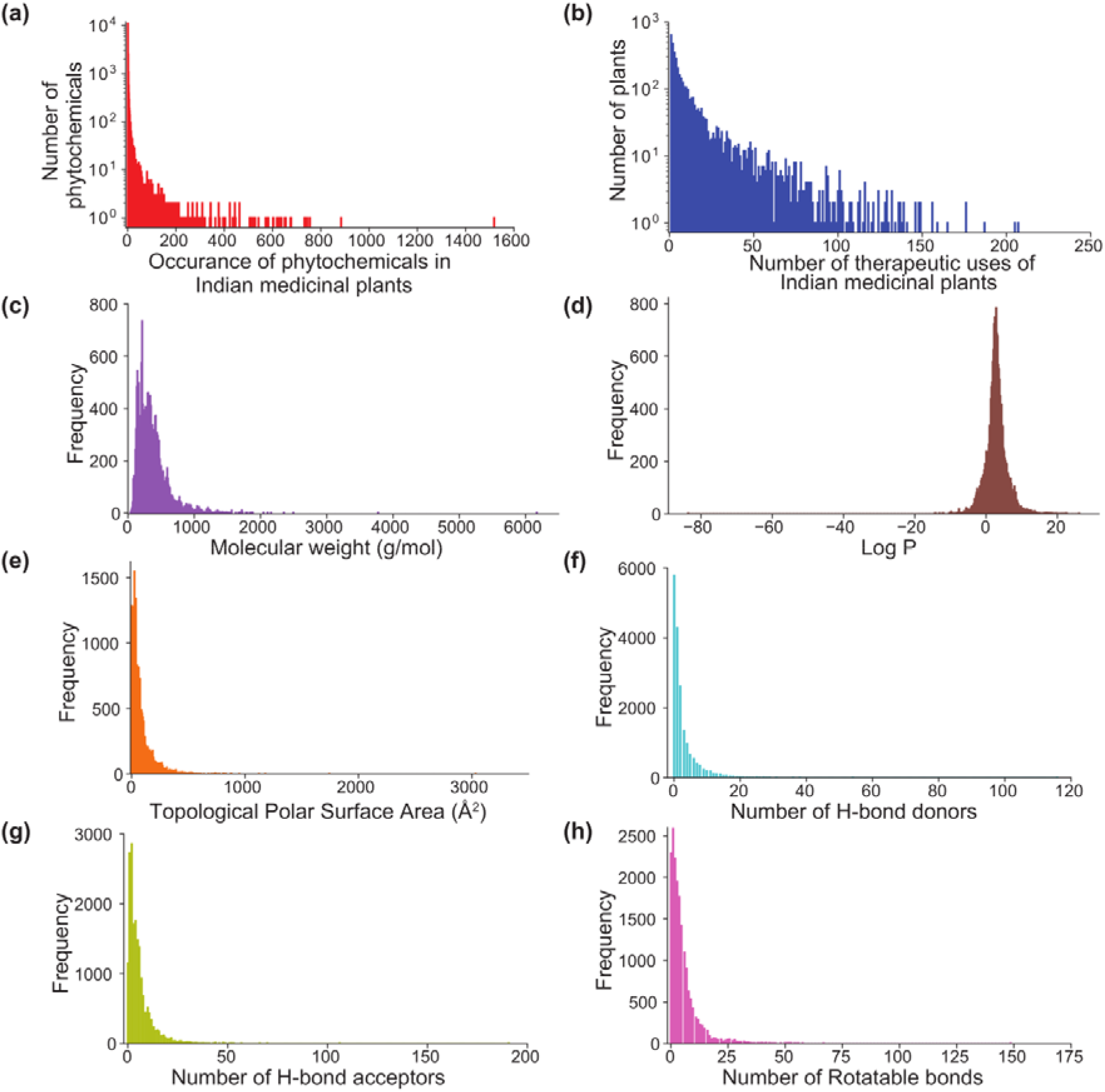
Basic statistics and distribution of the physicochemical properties for phytochemicals in IMPPAT 2.0. **(a)** Histogram of the number of Indian medicinal plants that produce a given phytochemical in IMPPAT 2.0. **(b)** Histogram of the number of therapeutic uses per Indian medicinal plant in IMPPAT 2.0. Distribution of six important physicochemical properties for 17967 phytochemicals, namely, **(c)** Molecular weight (g/mol), **(d)** log P, **(e)** Topological polar surface area (Å^2^), **(f)** number of hydrogen bond (H-bond) donors, **(g)** number of hydrogen bond (H-bond) acceptors, and **(h)** number of rotatable bonds.

### Enhanced annotation to enable exploration of the phytochemical space

We have significantly enhanced the additional information on phytochemicals in IMPPAT 2.0, and we now describe some of these new features in the updated database.

To make the phytochemical library of IMPPAT 2.0 compliant with Findable, Accessible, Interoperable, and Reusable (FAIR) principles, we assign unique IMPPAT identifiers to phytochemicals in the database, and thereafter, the identifiers are annotated with chemical names, structural features, and external links to standard chemical databases. Moreover, we provide the 2D and 3D chemical structures of phytochemicals in five different file formats (Methods).

Molecular scaffold represents the core structure of a molecule and is a key concept with wide applications in medicinal chemistry. In IMPPAT 2.0, we used the definition by Lipkus *et al.*^24,25^ to compute and provide the molecular scaffolds for phytochemicals at three levels (Methods). This scaffold information can be used by a chemist to group and retrieve phytochemicals with the same core structure to further build upon them. In IMPPAT 2.0, we also used the definition by Peter Ertl^26^ to provide the functional groups present in phytochemicals. This functional group information can also facilitate the exploration of the phytochemical space by chemists. Further enhancement of phytochemical annotation in IMPPAT 2.0 include new information such as DeepSMILES^27^ which is an adaptation of SMILES for use in machine learning, natural product specific chemical classification from NP classifier^28^, and natural product likeness (NP-likeness)^29^ score.

Molecular descriptors capture important structural features and are useful in machine learning based classification and regression analysis such as Quantitative Structure Activity Relationship (QSAR). In IMPPAT 2.0, we provide 1875 2D and 3D chemical descriptors for each phytochemical. Lastly, drug-likeness scores can enable selection of chemicals with favourable properties as drug lead molecules. In IMPPAT 2.0, we also evaluated the drug-likeness of phytochemicals based on multiple scores computed using in-house scripts (Methods).

Figure 3c-h shows the distribution of six important physicochemical properties for the 17967 phytochemicals in IMPPAT 2.0. Based on chemical classification obtained by ClassyFire^30^, the 17967 phytochemicals have been hierarchically categorized into 20 superclass, 250 class and 410 subclass. Among the 20 superclass, Lipids and lipid-like molecules, Phenylpropanoids and polyketides, and Organoheterocyclic compounds are the top three with 6904, 3007, and 2202 phytochemicals, respectively (Figure 4a). Further, using NP classifier^28^, the 17967 phytochemicals have been classified into one of seven biosynthetic pathways for natural products. Terpenoids, Shikimates and Phenylpropanoids, and Alkaloids are the top three biosynthetic pathways with 6049, 4206, and 2446 phytochemicals, respectively (Figure 4b).

**Figure 4:**
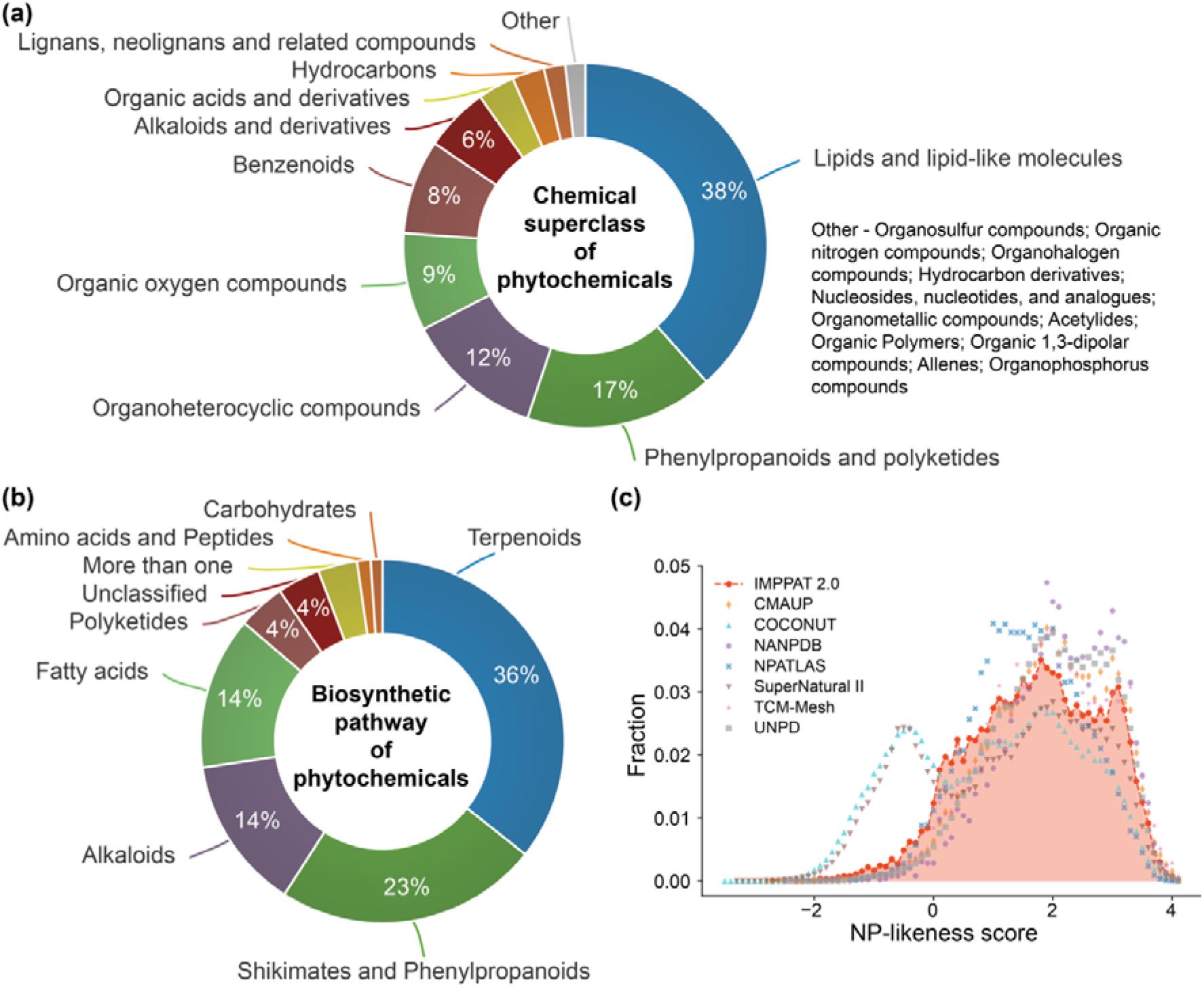
Chemical classification, biosynthetic pathways and natural product likeness of phytochemicals in IMPPAT 2.0. **(a)** Chemical superclass of phytochemicals predicted by ClassyFire^30^. **(b)** Biosynthetic pathways for phytochemicals predicted by NP classifier^28^. **(c)** Distribution of the NP-likeness scores for phytochemicals in IMPPAT 2.0 and other natural product libraries.

NP-likeness^29^ score is a measure to quantify the similarity of a given chemical structure to the natural product space. This score ranges from −5 to 5; the higher the score, more likely the molecule is a natural product^31^. Previous studies have shown that the NP-likeness of natural product libraries is predominantly positive, and moreover, is different from synthetic libraries which is predominantly negative^32,33^. On expected lines, phytochemicals in IMPPAT 2.0 have a predominantly positive NP-likeness score (>93%). Further, the distribution of the NP-likeness scores for phytochemicals in IMPPAT 2.0 is found to be similar to other natural product libraries (Figure 4c).

Lastly, IMPPAT 2.0 compiles information on 27365 predicted interactions between phytochemicals and human target proteins from STITCH^34^ database. These 27365 interactions involve 1294 phytochemicals and 5042 human target proteins.

### Increase in coverage of therapeutic uses

Building upon the compiled information in IMPPAT 1.0^10^, we enhanced the therapeutic use information in IMPPAT 2.0 to the level of plant parts and expanded to cover the 4010 Indian medicinal plants in the updated database. This information on therapeutic use of Indian medicinal plants was compiled from 146 books on traditional medicine (Supplementary Table S2). Only 9 out of these 146 books were covered in IMPPAT 1.0. Further, there are 56 books common to the set of 70 books from which phytochemical information was compiled and the set of 146 books from which therapeutic use information was compiled (Supplementary Tables S1-S2).

Since the therapeutic use of medicinal plants is reported using synonymous terms across different books, we undertook a manual curation effort to standardize the therapeutic use terms in IMPPAT 2.0. Specifically, we mapped the therapeutic use terms compiled from different books to standardized terms from Medical Subject Headings (MeSH; https://meshb.nlm.nih.gov/), International Classification of Diseases 11th Revision (ICD-11; https://icd.who.int/browse11/), Unified Medical Language System (UMLS; https://uts.nlm.nih.gov/uts/umls) and Disease Ontology (https://disease-ontology.org/). In the end, this effort to map the ethnopharmacological information on Indian medicinal plants to the standard vocabulary used in modern medicine led to a non-redundant list of 1095 standardized therapeutic use terms in IMPPAT 2.0.

Overall, there are 89733 non-redundant plant – part – therapeutic use associations in IMPPAT 2.0 spanning 4010 Indian medicinal plants and 1095 standardized therapeutic uses. At the level of plant – therapeutic use associations (after ignoring the plant parts), there is a 5-fold increase in IMPPAT 2.0 (Table 1). Figure 3b shows the histogram of the number of therapeutic uses per Indian medicinal plant in IMPPAT 2.0. While 21% of the Indian medicinal plants (851) in IMPPAT 2.0 have > 20 therapeutic uses, the majority of Indian medicinal plants (2488) have < 10 therapeutic uses.

### Increase in coverage of traditional medicinal formulations

Finally, IMPPAT 2.0 also compiles information on 7815 plant – part – traditional medicinal formulation associations which encompass 569 Indian medicinal plants and 1133 traditional Indian medicinal formulations (Table 1). This information was compiled using 1250 openly accessible formulations in Traditional Knowledge Digital Library (TKDL; http://www.tkdl.res.in) database. Further, the 1133 traditional Indian medicinal formulations in IMPPAT 2.0 belong to four systems of medicine, namely, Ayurveda (470), Unani (441), Siddha (187), and Sowa-Rigpa (35).

### Web design and data accessibility

The webserver for the previous version IMPPAT 1.0 enabled users to easily access the compiled information on Indian medicinal plants. Also, IMPPAT 1.0 webserver enabled cheminformatics analysis such as filtering phytochemicals based on their physicochemical properties, drug-likeness scores and chemical similarity. For the latest release, IMPPAT 2.0, we have completely redesigned the associated website: https://cb.imsc.res.in/imppat. While incorporating all the features of the previous version, the web-interface of IMPPAT 2.0 has multiple new features to facilitate the ease-of-use and exploration of the phytochemical space of Indian medicinal plants. This section describes some of the salient features of the IMPPAT 2.0 website. Users can access the compiled information in IMPPAT 2.0 via its web-interface by three means namely, *browse*, *basic search* and *advanced search*.

### Browse

In the web-interface, users can *browse* the compiled information in three different ways: (a) Phytochemical association, (b) Therapeutic use, and (c) Traditional medicinal formulation.

The phytochemical association section within browse enables a user to choose either an Indian medicinal plant, or a phytochemical, or a chemical superclass of phytochemicals, to retrieve compiled information in IMPPAT 2.0 on plant – part – phytochemical associations along with literature references. If a specific plant is chosen, the user is redirected to a new page containing plant-specific information along with a table listing the phytochemical constituents for the plant at the level of plant parts (Figure 5a). The page also displays a network visualization of the plant – phytochemical associations enabling the user to visually explore the phytochemical space of the chosen plant. If instead of choosing a specific plant in the phytochemical association section, the user chooses a phytochemical or a chemical superclass of phytochemicals, the user is redirected to a new page containing a table listing the plant – part – phytochemical associations for the chosen phytochemical or for the phytochemicals belonging to the chosen chemical superclass.

**Figure 5:**
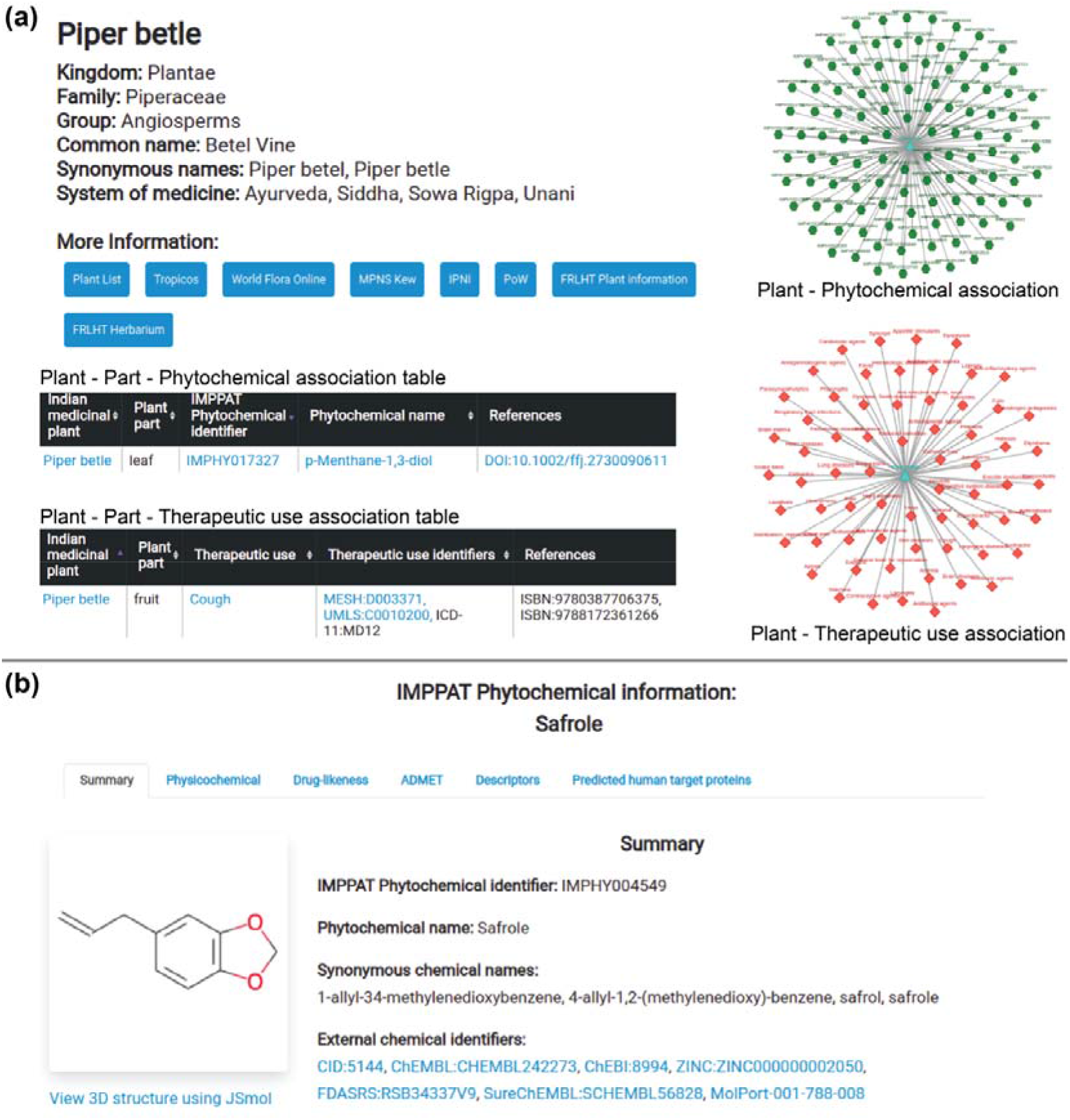
The web-interface of the IMPPAT 2.0 database. **(a)** Snapshots of the results of queries for a phytochemical or a therapeutic use of an Indian medicinal plant. In this example, we show from IMPPAT 2.0 for *Piper betle* the snapshots of the plant information, plant – part – phytochemical association table, plant – part – therapeutic use association table, and network visualization of plant – phytochemical associations and plant – therapeutic use associations. **(b)** Screenshot of the dedicated page containing detailed information for the phytochemical Safrole.

Similar to the phytochemical association section within browse, the therapeutic use association section enables users to retrieve compiled information in IMPPAT 2.0 on the plant – part – therapeutic use associations with literature references by choosing either an Indian medicinal plant, or a therapeutic use term (Figure 5a). The users can also retrieve compiled information in IMPPAT 2.0 on the plant – part – traditional medicinal formulation associations by choosing either an Indian medicinal plant, or a TKDL traditional medicinal formulation identifier, or a traditional Indian system of medicine such as Ayurveda, Siddha, Sowa-Rigpa, and Unani.

### Basic search

In the web-interface, users can perform text-based searches in the *basic search* section to retrieve compiled information. The basic search section has three tabs: (a) Phytochemical association, (b) Therapeutic use, and (c) Traditional medicinal formulation.

In the phytochemical association tab, a user can perform text-based search using complete or partial name of the plant, or IMPPAT phytochemical identifier, or complete or partial name of the phytochemical, to retrieve compiled information in IMPPAT 2.0 on plant – part – phytochemical associations. Upon submitting the text query, the user is presented with a table on the same page listing the relevant plant – part – phytochemical associations with literature references. In this table, the user can click any phytochemical name or identifier to view the page with detailed information on the phytochemical.

Similarly, in the therapeutic use tab, a user can perform text-based search using complete or partial name of the plant, or therapeutic use term, to retrieve compiled information in IMPPAT 2.0 on plant – part – therapeutic use associations with literature references as a table on the same page. Likewise, in the traditional medicinal formulation tab, a user can perform text-based search using complete or partial name of the plant, or TKDL formulation identifier, to retrieve compiled information in IMPPAT 2.0 on plant – part – traditional medicinal formulation associations as a table on the same page. In this table, on clicking the TKDL formulation identifier, the user is redirected to the corresponding formulation page in TKDL with additional information on the medicinal formulation.

### Advanced search

In the web-interface, the *advanced search* section enables a user to filter and retrieve a subset of phytochemicals compiled in IMPPAT 2.0 based on their physicochemical properties, drug-likeness, chemical similarity, and molecular scaffolds. The physicochemical filter tab provides a user with the option to retrieve phytochemicals of interest based on molecular weight, log P, topological polar surface area, hydrogen bond acceptors, hydrogen bond donors, number of heavy atoms, number of heteroatoms, number of rings, number of rotatable bonds, stereochemical complexity and shape complexity. Similarly, the drug-like filter tab enables a user to filter phytochemicals based on multiple drug-likeness scoring schemes.

The chemical similarity filter tab enables identification of phytochemicals in IMPPAT 2.0 that are structurally similar to a user submitted query compound. To submit a query compound, the user can either use the molecular editor to draw its chemical structure, and thereafter, search the corresponding SMILES, or directly enter the SMILES to perform the search. Upon submitting the SMILES of a query compound, the webserver will display a table listing the top 10 phytochemicals in IMPPAT 2.0 which are structurally similar based on Tanimoto coefficient (Tc)^35^, a standard measure to quantify the extent of chemical similarity (Methods). The scaffold filter tab enables a user to retrieve phytochemicals based on shared molecular scaffold. A user can select one of the three types of scaffold namely, graph/node/bond (G/N/B) level, or graph/node (G/N) level, or graph level (Methods), and thereafter, select the desired scaffold from the dropdown menu, to view the list of phytochemicals in the database having the desired scaffold. Overall, the advanced search page of IMPPAT 2.0 enables cheminformatics based exploration of the phytochemical space of the Indian medicinal plants towards natural product based drug discovery.

### Detailed information on phytochemicals

In the web-interface, a user is redirected to a dedicated page containing detailed information on a specific phytochemical upon clicking the corresponding phytochemical identifier or name in the tables fetched via *browse* or *basic search* or *advanced search* options. The dedicated page provides detailed information for a phytochemical in six tabs: (a) summary, (b) physicochemical, (c) drug-likeness, (d) ADMET, (e) descriptors, and (f) predicted human target proteins (Figure 5b). The summary tab provides basic information such as the chemical name, chemical classification, 2D and 3D chemical structures, molecular scaffolds, for the phytochemical. The remaining five tabs give the physicochemical properties, drug-likeness scores, predicted ADMET properties, molecular descriptors and predicted human target proteins from STITCH^34^ database, respectively, for the phytochemical. The predicted human target proteins tab also provides a network visualization of the phytochemical – predicted human target protein associations.

### Molecular complexity comparison with other collections of small molecules

Small molecules which are selective and specific binders of a target protein are preferable for drug development over promiscuous binders which can interact with both primary target and off-target proteins. Several molecular complexity metrics have been shown to correlate with the selectivity or promiscuity of small molecules^36,37^. In particular, Clemons *et al.*^38^ have shown that stereochemical complexity and shape complexity are excellent indicators of target protein specificity of small molecules.

In their work, Clemons *et al.*^38^ correlated the distribution of stereochemical and shape complexity with protein binding specificity of three different representative small molecule collections namely, commercial compounds (CC), diversity-oriented synthesis compounds (DC’) and natural products (NP) (Methods). Clemons *et al.*^38^ found that CC, DC’ and NP molecules on an average have low, intermediate and high values, respectively of both stereochemical and shape complexity. Thereafter, Clemons *et al.*^38^ correlated the two molecular complexities to protein binding specificities to find that CC molecules with low complexity are enriched in promiscuous binders and depleted in specific binders, while in comparison DC’ molecules with intermediate complexity and NP molecules with high complexity are more enriched in specific binders and depleted in promiscuous binders. Lastly, NP molecules were found to be more depleted in promiscuous binders in comparison to DC’ molecules^38^.

Previously^10^, we compared the stereochemical and shape complexity of the CC, DC’ and NP molecules with 9596 phytochemicals in IMPPAT 1.0 from Indian medicinal plants and 10140 phytochemicals in TCM-Mesh^39^ from Chinese medicinal plants. In a nutshell, we showed conclusively that phytochemicals in both IMPPAT 1.0 and TCM-Mesh are similar to NP collection in terms of their distributions of stereochemical and shape complexity. Due to significant increase in the number of phytochemicals in IMPPAT 2.0, we compared the distribution of stereochemical and shape complexity of CC, DC’ and NP molecules with phytochemicals in IMPPAT 1.0 and IMPPAT 2.0 (Figure 6a). We find that the distributions of stereochemical and shape complexity for phytochemicals in IMPPAT 2.0 are very similar to IMPPAT 1.0, and closer to NP rather than DC’ or CC collections (Figure 6a).

**Figure 6:**
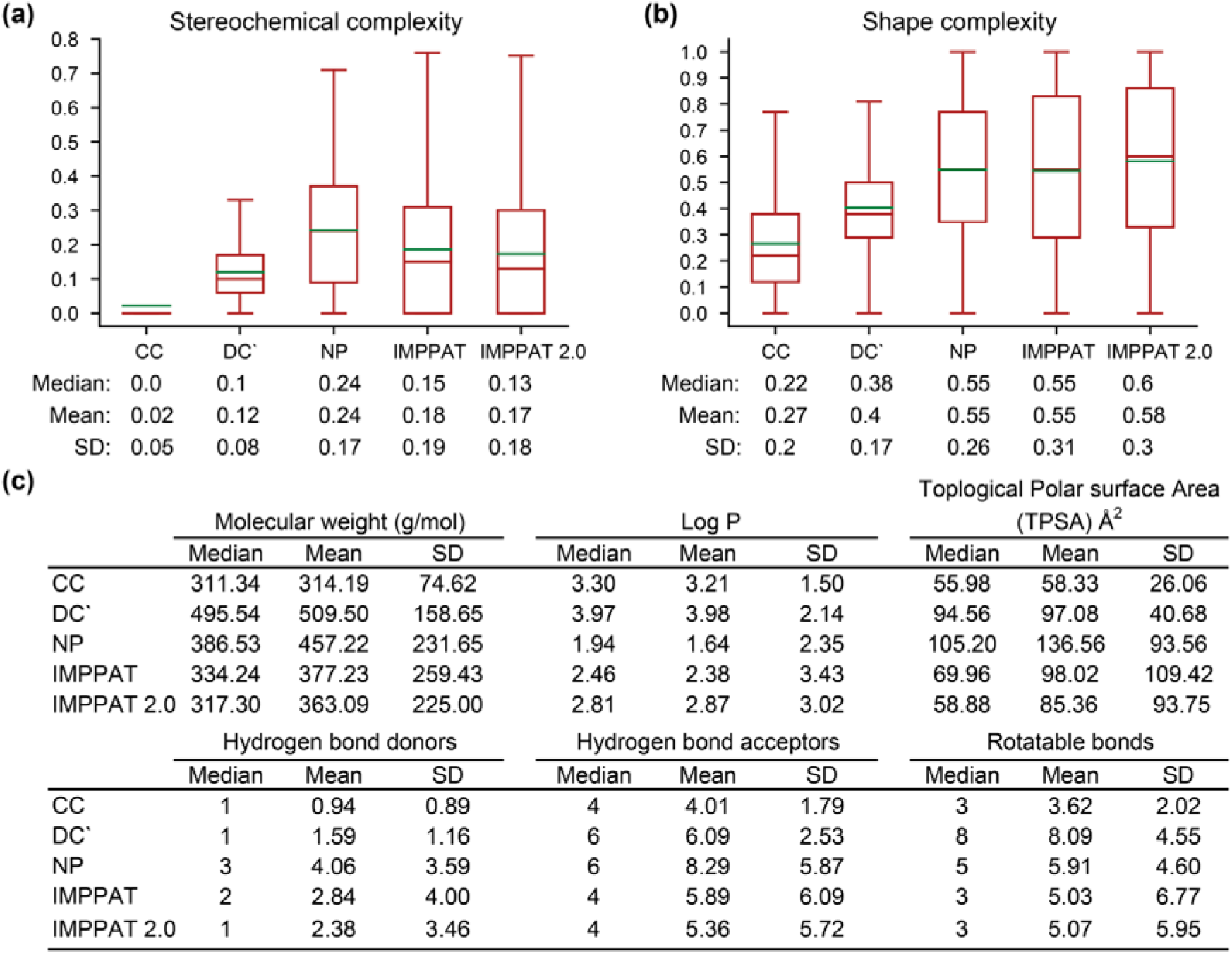
Comparison of the molecular complexity of chemical libraries. **(a)** Distribution of the stereochemical complexity, and **(b)** the shape complexity for small molecules in five chemical libraries, namely, CC, DC’, NP, IMPPAT version 1.0 and IMPPAT 2.0. Note that the lower end of the box plot is the first quartile, upper end is the third quartile, brown line inside the box is the median, green line is the mean of the distribution. Also, the median, mean and standard deviation (SD) of the distribution is shown below the box plot. **(c)** Median, mean and SD for six physicochemical properties, namely, Molecular weight (g/mol), log P, topological polar surface area (TPSA) (Å^2^), number of hydrogen bond donors, number of hydrogen bond acceptors, and number of rotatable bonds, for small molecules in five chemical libraries.

In another study, Clemons *et al.*^40^ have shown that CC, DC’ and NP occupy different regions in the physicochemical space defined by six properties namely, molecular weight, log P, topological polar surface area, number of hydrogen bond donors, number of hydrogen bond acceptors, and number of rotatable bonds. In terms of these six physicochemical properties, we also find that phytochemicals in IMPPAT 2.0 are very similar to IMPPAT 1.0, and closer to NP and DC’ rather than CC collection (Figure 6b).

Overall, our analysis of the molecular complexities of the phytochemicals in IMPPAT 2.0 finds that the phytochemical space of Indian medicinal plants has many similarities with other natural product spaces. Notably, the phytochemical space is likely to be enriched in specific protein binders, and therefore, a valuable space for ongoing efforts in drug discovery.

### Molecular scaffold based structural diversity

Analysis of the structural diversity of a chemical space has significance for the discovery of new and novel small molecule entities. The concept of molecular scaffolds has emerged as one of the reliable ways to quantify the structural diversity^41^ of chemical libraries. One way to define the molecular scaffold is via the core structure of a molecule with all its ring system and all chain fragments connecting the rings^41,42^. Previously, Lipkus *et al.*^24,25^ have analyzed the scaffold diversity of organic compounds compiled in Chemical Abstracts Service (CAS) database to find that the frequency distribution of scaffolds is uneven, with most scaffolds occurring in a small number of molecules and few scaffolds occurring in a very large number of molecules. To quantify the scaffold diversity of the phytochemicals in IMPPAT 2.0, we followed Lipkus *et al.*^24,25^ to compute the molecular scaffold at three levels, namely, graph/node/bond (G/N/B) level, graph/node (G/N) level and graph level (Methods). Among the phytochemicals in IMPPAT 2.0, we find 5179 scaffolds at G/N/B level, 4072 at G/N level and 3434 at graph level.

Thereafter, we compared the scaffold diversity of IMPPAT 2.0 with seven other natural product libraries (CMAUP^43^, COCONUT^44^, NANPDB^45^, NPATLAS^46^, SuperNatural-II^47^, TCM-Mesh^39^ and UNPD), approved drugs obtained from Drugbank^48^, and more than 100 million organic compounds from PubChem^23^ (Table 2). Focusing solely on scaffolds at the G/N/B level, we find that phytochemical space of IMPPAT 2.0 is the third highest among the seven natural product libraries in terms of the fraction of scaffolds per molecule (N/M) and the fraction of singleton scaffolds per molecule (N_sing_/M), after TCM-Mesh and NANPDB (Table 2).

**Table 2:** Scaffold diversity of phytochemicals in IMPPAT 2.0, and comparison with other chemical libraries. The molecular scaffolds are computed at graph/node/bond (G/N/B) level. Here, M is number of molecules with scaffold and this number is less than the library size as linear molecules with no ring system have no scaffolds. Further, N is the number of scaffolds, N_sing_ is the number of singleton scaffolds, AUC is the area under the curve, and P_50_ is the percentage of scaffolds that account for 50% of the chemical library.

| Chemical library | M | N | $N_{\text{sing}}$ | N/M | $N_{\text{sing}}/M$ | $N_{\text{sing}}/N$ | AUC | $P_{50}$ |
| --- | --- | --- | --- | --- | --- | --- | --- | --- |
| Approved drugs | 2097 | 1255 | 1012 | 0.6 | 0.48 | 0.81 | 0.69 | 17.93 |
| TCM-Mesh | 9417 | 3946 | 2626 | 0.42 | 0.28 | 0.67 | 0.75 | 11.02 |
| NANPDB | 4645 | 1762 | 1093 | 0.38 | 0.24 | 0.62 | 0.76 | 10.67 |
| IMPPAT 2.0 | 15226 | 5179 | 3338 | 0.34 | 0.22 | 0.64 | 0.79 | 6.58 |
| NPATLAS | 31099 | 10227 | 5947 | 0.33 | 0.19 | 0.58 | 0.79 | 8.35 |
| COCONUT | 385926 | 109024 | 65963 | 0.28 | 0.17 | 0.61 | 0.82 | 4.82 |
| CMAUP | 43987 | 11105 | 6151 | 0.25 | 0.14 | 0.55 | 0.82 | 5.15 |
| UNPD | 215585 | 44281 | 22514 | 0.21 | 0.1 | 0.51 | 0.85 | 3.39 |
| SuperNatural II | 308998 | 62125 | 30453 | 0.2 | 0.1 | 0.49 | 0.85 | 3.61 |
| PubChem | 101452728 | 12493379 | 7059386 | 0.12 | 0.07 | 0.57 | 0.91 | 0.22 |

Figure 7a,b show the distribution of the number of rings and number of heteroatoms across the 5179 scaffolds at G/N/B level found in phytochemicals of IMPPAT 2.0. While more than 74% of the 5179 scaffolds are relatively small with ≤ 5 rings in them, only 2.5% of the scaffolds have ≥ 10 rings (Figure 7a). Notably, 231 scaffolds (4.5%) are single ring system, and this indicates high degree of ring diversity in phytochemicals of IMPPAT 2.0. We also find 49.7% of the 5179 scaffolds have two or three or four heteroatoms, and only 0.4% of the scaffolds contain ≥ 20 heteroatoms (Figure 7b). Further, 518 scaffolds (10%) are completely composed of carbon atoms. Figure 7a,b also show that the distributions of number of rings and number of heteroatoms in scaffolds found in phytochemicals of IMPPAT 2.0 are similar to respective distributions for other natural product libraries, approved drugs, and organic compounds from PubChem.

**Figure 7:**
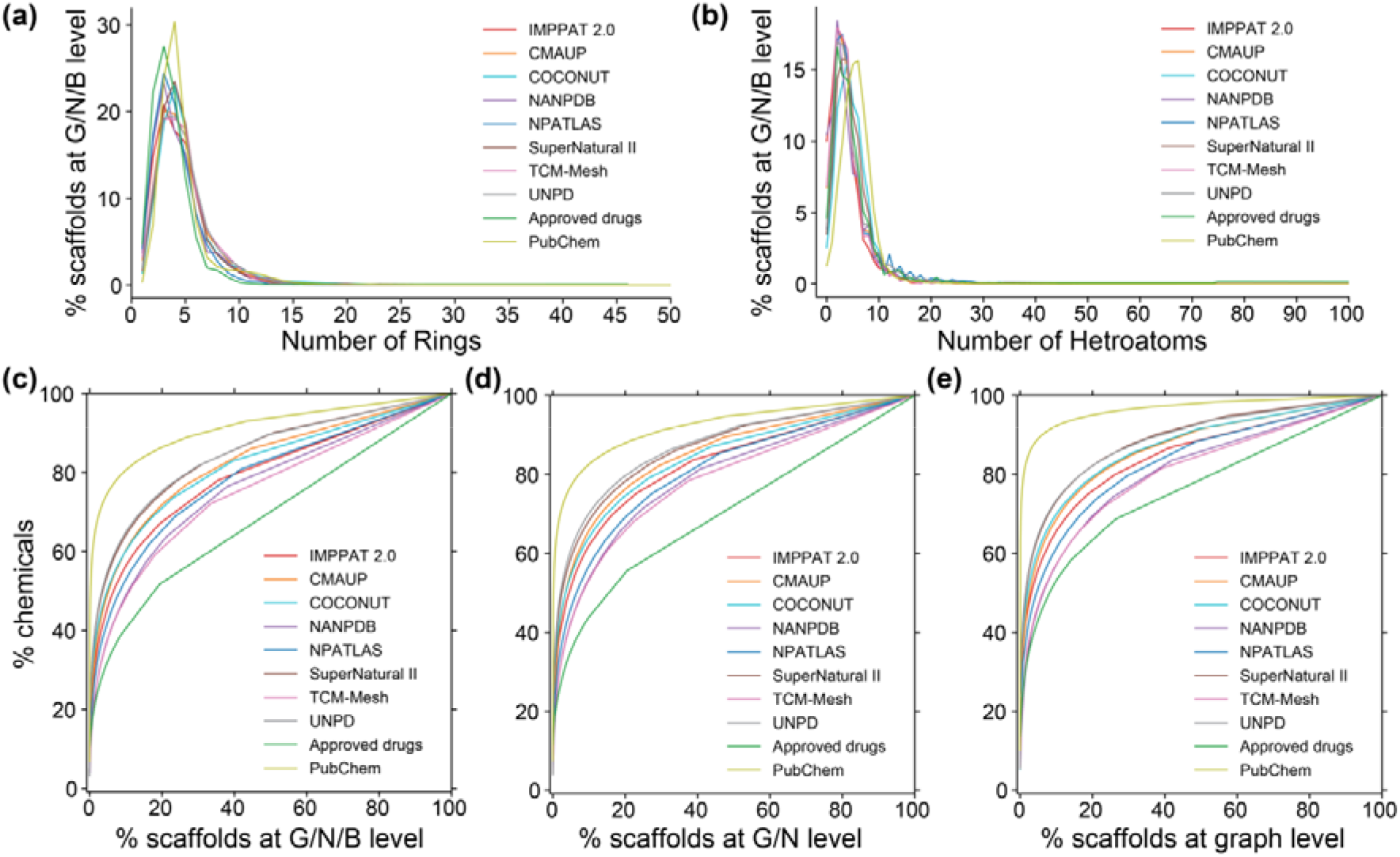
Analysis of the scaffold diversity of phytochemicals in IMPPAT 2.0 with seven other natural product libraries, approved drugs, and organic compounds from PubChem^23^. Distribution of **(a)** the number of ring systems and **(b)** the number of heteroatoms, in scaffolds at graph/node/bond (G/N/B) level. Cyclic system retrieval (CSR) curves for scaffolds at: **(c)** G/N/B level, **(d)** graph/node (G/N) level, and **(e)** graph level.

To further understand and compare the structural diversity of the phytochemical space of IMPPAT 2.0 with other chemical libraries, cyclic system retrieval (CSR) curves^24,25,49,50^ were plotted for scaffolds computed at G/N/B level (Figure 7c), G/N level (Figure 7d) and graph level (Figure 7e). CSR curves were generated by plotting the percent of scaffolds on the x-axis and the percent of compounds that contain those scaffolds on the y-axis. From the CSR curves, metrics such as area under the curve (AUC) and percent scaffolds required to retrieve 50% of the compounds (P_50_) were computed. Notably, several studies have used the above metrics to quantify and compare scaffold diversity of chemical libraries^24,25,49–51^. In an ideal distribution with maximum scaffold diversity wherein each compound has a unique scaffold, the CSR curve will be the diagonal line with AUC value of 0.5. It is seen that the CSR curves for phytochemicals in IMPPAT 2.0 (red) and other chemical libraries rise steeply and then levels off (Figures 7c-e). As we move from scaffolds at G/N/B level (least abstraction) to G/N level to graph level (high abstraction), the scaffold diversity reduces across all the chemical libraries, with CSR curves shifting up away from the diagonal (Figures 7c-e).

Importantly, the scaffold diversity of phytochemicals in IMPPAT 2.0 (red) and other natural product libraries lie in between the scaffold diversity of 100 million organic compounds from PubChem (low diversity) and approved drugs (high diversity) (Figure 7c-e). Table 2 lists the AUC and P_50_ from CSR curves of scaffolds at G/N/B level for the phytochemicals in IMPPAT 2.0 and other chemical libraries. In line with expectation, the approved drug library was found to be most diverse with AUC of 0.69 and P_50_ of 17.93% (Table 2). Interestingly, the scaffold diversity of phytochemicals in IMPPAT 2.0 was found to be greater than the entire organic compound library from PubChem, and moreover, it is the third or fourth most diverse library among the eight natural product libraries based on AUC of 0.79 and P_50_ of 6.58%, respectively (Table 2). Further, 64.5% of the 5179 scaffolds at G/N/B level found in phytochemicals of IMPPAT 2.0 are singletons which are present in only one compound (Table 2). In contrast, 217 scaffolds present in 10 or more phytochemicals cumulatively account for 43.6% of the phytochemicals in IMPPAT 2.0, and a molecule cloud visualization^52,53^ of these scaffolds is shown in Figure 8 (after excluding benzene ring scaffold). In sum, these results highlight that the phytochemical space of IMPPAT 2.0 is structurally diverse with high scaffold diversity in comparison with the organic compounds from PubChem, and moreover, has similar scaffold diversity as other large natural product libraries.

**Figure 8:**
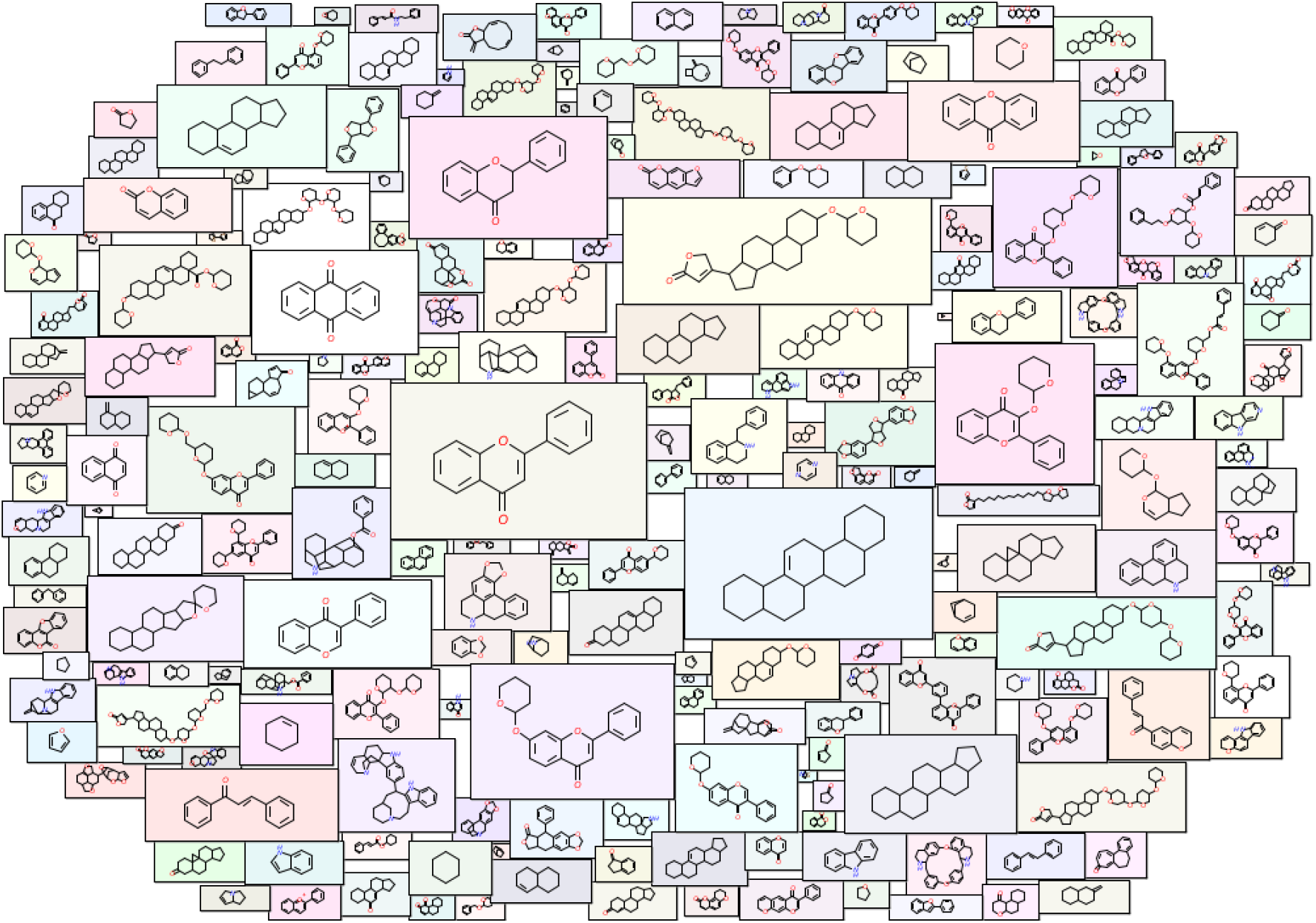
Molecular cloud visualization^52,53^ of the top scaffolds at G/N/B level present in phytochemicals of IMPPAT 2.0. The top constitute the 217 scaffolds at G/N/B level that are present in ≥ 10 phytochemicals in IMPPAT 2.0. In this figure, 216 of these top scaffolds are shown after excluding the benzene ring (which is the most frequent scaffold in all large chemical libraries). Here, the size of the structure is proportional to the frequency of occurrence of the scaffold in phytochemicals of IMPPAT 2.0.

### Drug-like phytochemical space

Natural products have been an important source of approved drugs^8,54^. To predict the subset of drug-like phytochemicals in IMPPAT 2.0, we used six scoring schemes namely, Lipinski’s rule of five (RO5)^55^, Ghose rule^56^, Veber rule^57^, Egan rule^58^, Pfizer 3/75 rule^59^ and GlaxoSmithKline’s (GSK) 4/400 rule^60^. Figure 9a is an UpSet^61^ visualization of the set intersections of phytochemicals that pass one or more of these six rules. Majority of the phytochemicals pass RO5 (14847), followed by Veber (13574) and Egan (12390) rules. Pfizer 3/75 was found to be the most restrictive rule, with 4924 phytochemicals passing it. A drug-like subset of 1335 phytochemicals is identified based on the stringent criteria of passing all six rules (Figure 9a; Supplementary Table S3).

**Figure 9:**
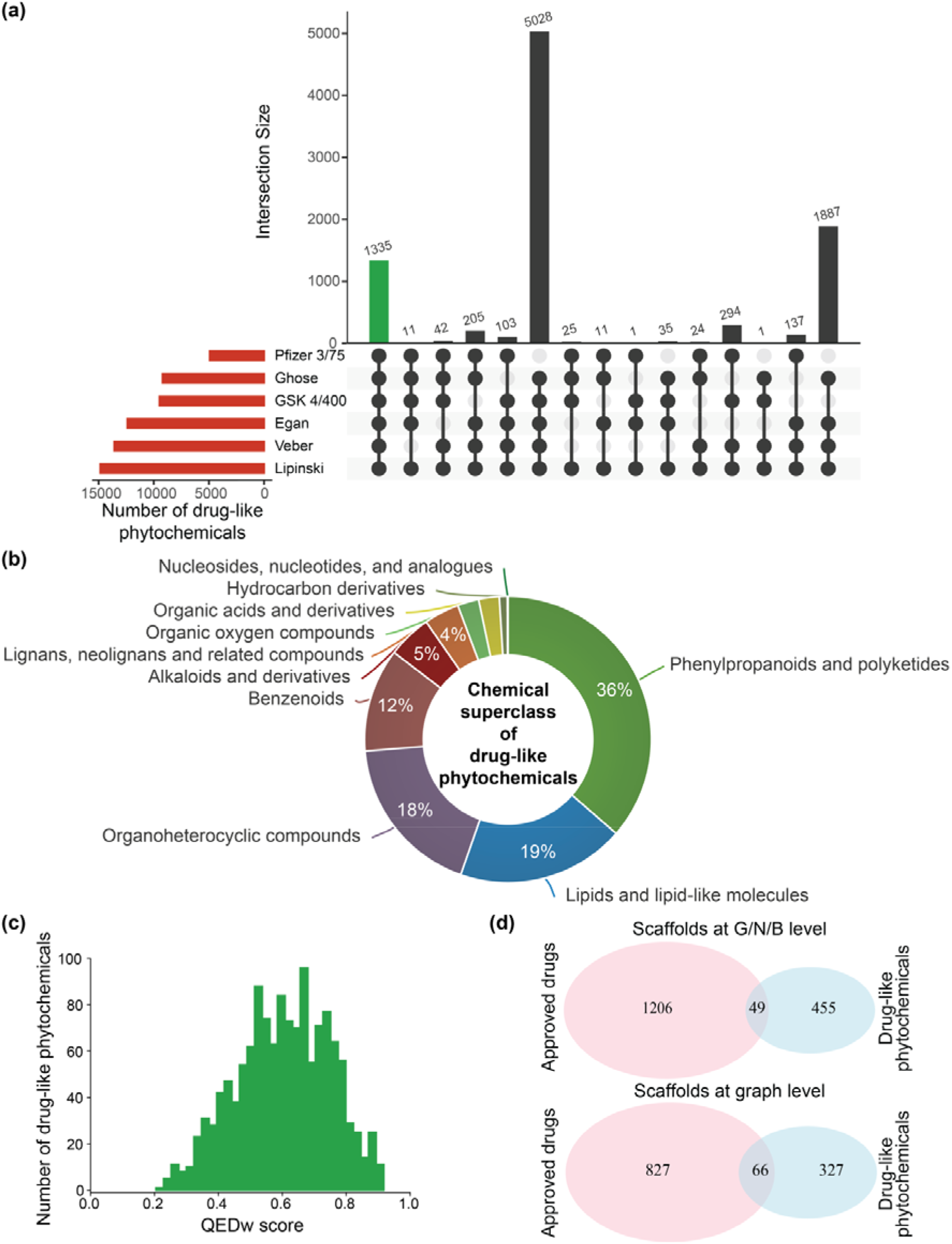
Drug-likeness analysis of phytochemicals in IMPPAT 2.0. **(a)** UpSet plot visualization of the set intersections of phytochemicals that pass one or more of the six drug-likeness rules. The horizontal bars show the number of phytochemicals which pass the different drug-likeness rules. The vertical bars show the set intersections between phytochemicals that pass different drug-likeness rules. The green bar shows the 1335 phytochemicals which pass all six drug-likeness rules. This plot was generated using UpSetR package^61^. **(b)** Chemical superclass of the 1335 drug-like phytochemicals as predicted by ClassyFire. **(c)** Distribution of QEDw scores for the 1335 drug-like phytochemicals. **(d)** Common scaffolds at the graph/node/bond (G/N/B) level and the graph level between the space of 1335 drug-like phytochemicals and approved drugs.

The top 5 plants in IMPPAT 2.0 based on associated drug-like phytochemicals are *Senna obtusifolia* (22), *Artemisia annua* (21), *Ailanthus altissima* (19), *Catharanthus roseus* (19) and *Senna tora* (19). Figure 9b shows the chemical classification for the 1335 drug-like phytochemicals obtained using ClassyFire^30^. The top 3 chemical superclasses namely, Phenylpropanoids and polyketides, Lipids and lipid-like molecules, and Organoheterocyclic compounds account for 486, 253, and 245 drug-like phytochemicals, respectively.

Weighted quantitative estimate of drug-likeness (QEDw) score can also be used to assess drug-likeness of small molecules, and this measure can take values between 0 (least drug-like) to 1 (most drug-like)^62^. For the 1335 drug-like phytochemicals, Figure 9c shows the distribution of QEDw scores with a mean of 0.60 and a standard deviation of 0.14. Notably, 104 of the drug-like phytochemicals have a high QEDw score ≥ 0.80.

We also compared the 1335 drug-like phytochemicals in IMPPAT 2.0 with the drugs approved by United States Federal Drug Administration (US FDA). A set of 2567 approved drugs were obtained from DrugBank^48^ version 5.1.9. Based on chemical similarity (Tc ≥ 0.50; Methods), we find 130 drug-like phytochemicals to be similar to one or more approved drugs. Interestingly, 11 drug-like phytochemicals in IMPPAT 2.0 are already US FDA approved drugs.

To assess the overlap in core chemical structure, we next computed the molecular scaffolds for the 1335 drug-like phytochemicals and 2567 approved drugs. At the G/N/B, G/N and graph levels, the 1335 drug-like phytochemicals were found to have 504, 444 and 393 scaffolds, respectively, while the 2567 approved drugs have 1255, 1171 and 893 scaffolds, respectively. Importantly, the drug-like phytochemicals and approved drugs share only 49, 60 and 66 scaffolds at G/N/B, G/N and graph levels, respectively (Figure 9d). Thus, the drug-like phytochemicals in IMPPAT 2.0 presents a unique chemical scaffold space with minimal overlap with approved drugs. These results highlight the potential of our database in aiding the ongoing hunt for new bioactive molecules.

By constructing a chemical similarity network (CSN), we next analyzed the structural diversity of the drug-like space of 1335 phytochemicals (Methods). Figure 10a shows the drug-like CSN wherein nodes correspond to phytochemicals and an edge exists between any pair of phytochemicals if Tc ≥ 0.5. The drug-like CSN is very sparse with graph density of 0.01, and it can be partitioned into 90 connected components (with at least 2 nodes each) and 210 isolated nodes. In Figure 10a, the top 12 connected components in terms of the number of constituent nodes are labeled. For instance, the connected component labeled 9 consists of 16 phytochemicals of which 2 phytochemicals (Colchicine and its metabolite Colchiceine) are approved drugs and remaining phytochemicals are similar to them. For each of the top 12 components, the maximum common substructure (MCS) is shown in Figure 10b; the substructures confirm the structural uniqueness of the different connected components (Methods). In sum, the CSN highlights the chemical dissimilarity, and hence, the structural diversity of the drug-like space of 1335 phytochemicals.

**Figure 10:**
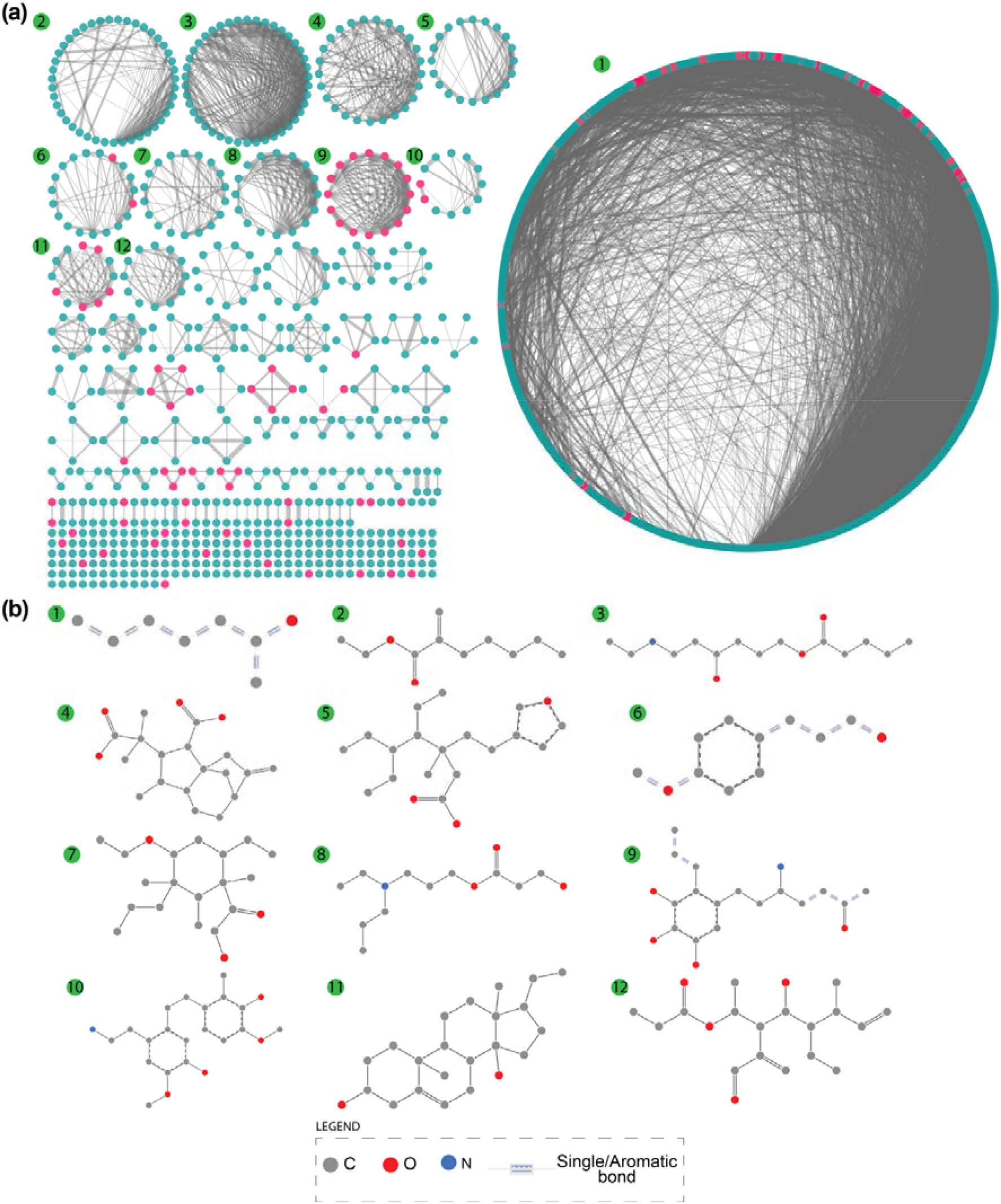
**(a)** Chemical similarity network (CSN) of the 1335 drug-like phytochemicals in IMPPAT 2.0. The degree sorted circle layout in Cytoscape^75^ is used to visualize the CSN. Cyan nodes correspond to drug-like phytochemicals that are not similar to any approved drug and pink nodes to those that are similar to at least one approved drugs. Edge thickness is proportional to the chemical similarity between the pair of drug-like phytochemicals. **(b)** Visualization of the SMARTS corresponding to the maximum common substructure (MCS) for the top 12 connected components obtained using SMARTSview webserver^72,73^.

### Comparison with the phytochemical space of Chinese medicinal plants

Previously^10^, a comparison of the 9596 phytochemicals in IMPPAT 1.0 with the 10140 phytochemicals in TCM-Mesh^39^ revealed that less than 25% phytochemicals (2305) in IMPPAT 1.0 are present in the TCM-Mesh. Notably, TCM-Mesh is a large-scale database compiling information on 10140 phytochemicals produced by 6235 Chinese medicinal plants^39^. We also performed a comparison of the 17967 phytochemicals in IMPPAT 2.0 with the 10140 phytochemicals in TCM-Mesh. Though the number of phytochemicals common to IMPPAT 2.0 and TCM-Mesh has increased to 3342, the percentage of the phytochemical space of IMPPAT 2.0 which is shared with TCM-Mesh has decreased to 18.6% (Figure 11a).

**Figure 11:**
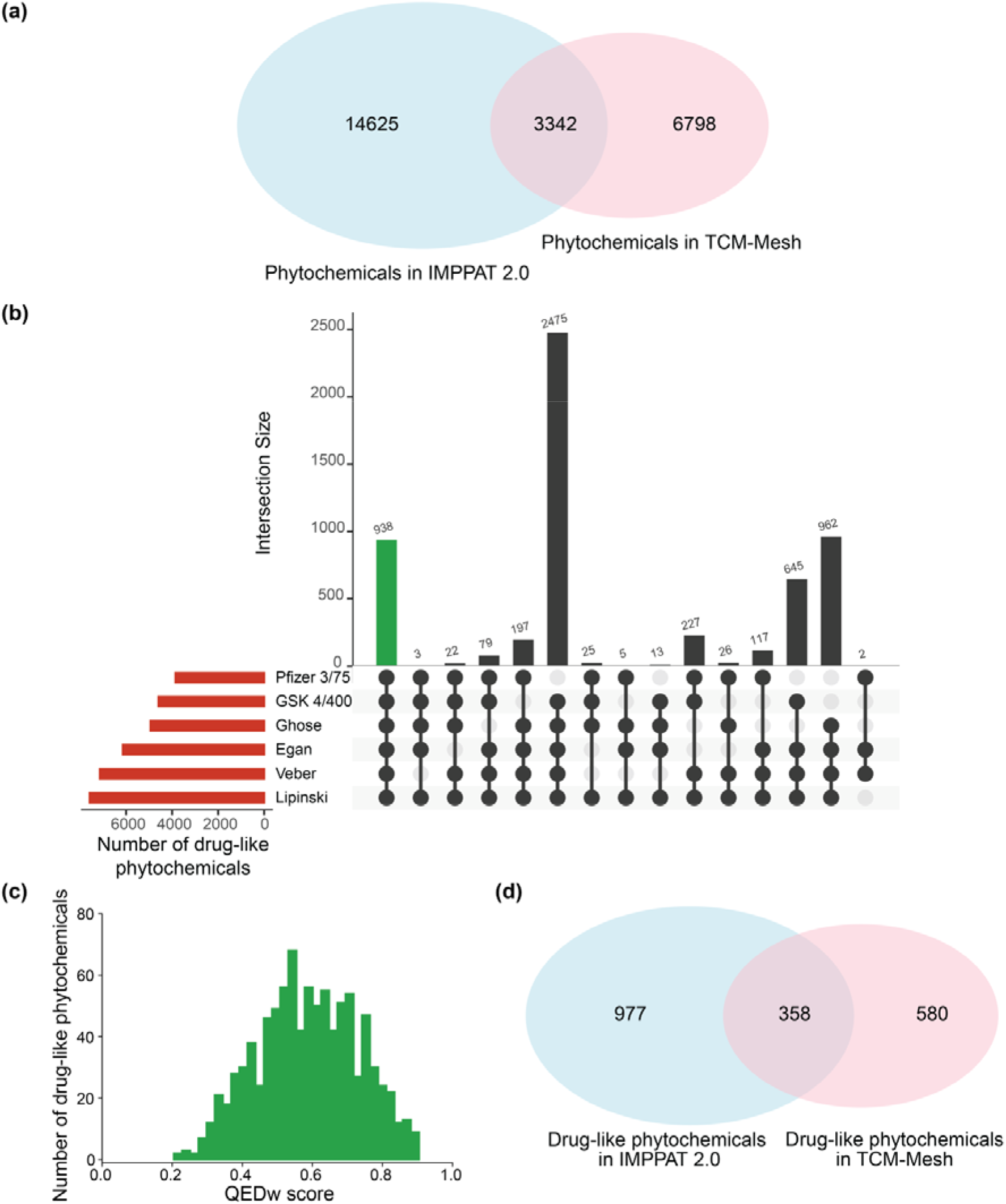
Comparison of the phytochemical space of Indian medicinal plants and Chinese medicinal plants. **(a)** Venn diagram shows the overlap between the phytochemicals in IMPPAT 2.0 and TCM-Mesh. **(b)** UpSet plot visualization of the set intersections of phytochemicals in TCM-Mesh that pass one or more of the six drug-likeness rules. The horizontal bars show the number of phytochemicals which pass the different drug-likeness rules. The vertical bars show the set intersections between phytochemicals that pass different drug-likeness rules. The green bar shows the 938 phytochemicals which pass all six drug-likeness rules. **(c)** Distribution of QEDw scores for the 938 drug-like phytochemicals in TCM-Mesh. **(d)** Venn diagram shows the overlap between the drug-like phytochemicals in IMPPAT 2.0 and TCM-Mesh.

Further, we compared the drug-like subset of 1335 phytochemicals in IMPPAT 2.0 with the corresponding drug-like subset in TCM-Mesh (Methods). Specifically, a subset of 938 drug-like phytochemicals was obtained in TCM-Mesh based on the six rules (Figure 11b). Further, Figure 11c shows the distribution of QEDw scores for the 938 drug-like phytochemicals in TCM-Mesh, and this distribution has a mean value of 0.59 and standard deviation of 0.14, similar to the distribution for the 1335 drug-like phytochemicals in IMPPAT 2.0. Lastly, there is a minor overlap of 338 phytochemicals between the subsets of drug-like phytochemicals in IMPPAT 2.0 and TCM-Mesh. These analyses attest to the uniqueness of the phytochemical spaces of Indian herbs and Chinese herbs, and therefore, the phytochemical atlas IMPPAT 2.0 is expected to further enrich the space of natural products.

## Discussion

In this contribution, we present IMPPAT 2.0, an enhanced and expanded database, compiling information via extensive manual curation on Indian medicinal plants, their phytochemicals, therapeutic uses and traditional medicine formulations. IMPPAT 2.0 is by far the largest phytochemical atlas specific to Indian medicinal plants to date.

In the updated database, we have more than doubled the coverage of Indian medicinal plants and nearly doubled the size of the phytochemical space. Further, we compile the phytochemicals, therapeutic uses and traditional medicinal formulations of the Indian medicinal plants at the level of plant parts. At the level of associations, IMPPAT 2.0 compiles 189386 plant – part – phytochemical, 89733 plant – part – therapeutic use, and 7815 plant – part – traditional medicinal formulation associations. Importantly, IMPPAT 2.0 provides a FAIR^18^ compliant non-redundant *in silico* stereo-aware library of 17967 phytochemicals. The phytochemical library has been annotated with several features including 2D and 3D chemical structures, molecular scaffolds, predicted human target proteins, physicochemical properties, drug-likeness scores and predicted ADMET properties. This will enable the effective use of the phytochemical library for screening efforts towards drug discovery. Also, the 1095 standardized therapeutic use terms in IMPPAT 2.0 are mapped to standard terms such as MeSH (https://meshb.nlm.nih.gov/), ICD-11 (https://icd.who.int/browse11/), UMLS (https://uts.nlm.nih.gov/uts/umls) and Disease Ontology (https://disease-ontology.org/) used in western medicine. Further, IMPPAT 2.0 web-interface has been completely redesigned to facilitate ease of use and to serve as a cheminformatics platform for exploring the phytochemical space of Indian medicinal plants. For instance, the advanced search page now enables the user to draw the chemical structure using a visual molecular editor to search for similar phytochemicals in the database and also allows the user to select the phytochemicals based on molecular scaffolds.

The cheminformatics analysis of the phytochemicals in IMPPAT 2.0 revealed that their stereochemical complexity and shape complexity is similar to the other natural products. Our analysis suggests that, like the library in IMPPAT 1.0, the phytochemicals in IMPPAT 2.0 are also more likely to be enriched with specific protein binders rather than promiscuous binders. The structural diversity analysis using molecular scaffolds has shown that the phytochemicals in IMPPAT 2.0 are structurally diverse with scaffold diversity similar to large natural product databases. Also, we find that the scaffold diversity of natural product libraries including IMPPAT 2.0 lies in between the scaffold diversity of more than 100 million organic compounds from PubChem (low diversity) and approved drugs (high diversity). This highlights the utility of our phytochemical library for the identification of biologically active new chemical entities with novel scaffolds. Using six drug-likeness scores, we identified a subset of 1335 drug-like phytochemicals which pass all six rules considered here. We find that only 11 of the drug-like phytochemicals are already approved drugs. Also, the drug-like phytochemicals and approved drugs have very few common scaffolds, revealing the pool of scaffolds present in drug-like phytochemicals in IMPPAT 2.0 but not present in approved drugs. Further, the chemical similarity network of the drug-like phytochemicals highlights the structural diversity of the drug-like space in IMPPAT 2.0. Finally, the comparison with the phytochemicals from Chinese medicinal plants shows that there is minimal overlap with the phytochemicals from Indian medicinal plants compiled in IMPPAT 2.0. These results show the uniqueness of the phytochemical space of IMPPAT 2.0 and its potential to further enrich the natural product chemical space.

In conclusion, IMPPAT 2.0 is a unique database enabling computational and experimental research in the area of natural product and traditional knowledge based drug discovery. In future, we will continue to expand, enhance and develop this unique platform to explore the phytochemical space of Indian medicinal plants.

## Methods

### Plant annotation

For the 4010 Indian medicinal plants in IMPPAT 2.0, the taxonomic information on kingdom, family and group was compiled using The Plant List database (http://www.theplantlist.org/). The common names of the Indian medicinal plants were obtained from the Flowers of India database (http://www.flowersofindia.net/), which compiles information for more than 6000 Indian plants. The IUCN Red List of Threatened species^19^ (https://www.iucnredlist.org/) is the most comprehensive resource on global conservation status of animals, fungi and plant species, and this list was used to ascertain the extinction risk of Indian medicinal plants. The usage of Indian medicinal plants in different traditional Indian systems of medicine such as Ayurveda, Siddha, Unani, Sowa-Rigpa and Homeopathy was manually compiled from pharmacopoeias published by Government of India and Traditional Knowledge Digital Library (TKDL; http://www.tkdl.res.in) of the Council of Scientific and Industrial Research, Government of India.

For the Indian medicinal plants in IMPPAT 2.0, we provide cross-reference links to associated information in other standard databases such as The Plant List, Tropicos (https://www.tropicos.org/), Encyclopedia of Indian medicinal plants from FRLHT (http://envis.frlht.org/), Medicinal Plants Names Service (MPNS; https://mpns.science.kew.org/), International Plant Names Index (IPNI; https://www.ipni.org/), Plants of the World Online (POW; https://powo.science.kew.org/), World Flora Online (WFO; http://www.worldfloraonline.org/) and Gardeners’ World (https://www.gardenersworld.com/).

### Phytochemical information

The 2D chemical structures of phytochemicals were converted to SDF, MOL and MOL2 file formats using OpenBabel^63^. The images of the 2D structures of phytochemicals were generated using RDKit^64^. The 3D chemical structures of phytochemicals were retrieved from PubChem^23^. If the 3D structure for a phytochemical was not available in PubChem, the 3D structure was generated from its 2D structure using RDKit by first embedding the 2D structure using ETKDG method and thereafter energy minimizing the structure using MMFF94 force field^64^. The 3D structures of phytochemicals were converted to SDF, MOL, MOL2, PDB and PDBQT file formats using OpenBabel^63^. Note that IMPPAT 2.0 provides 3D structures for 17910 phytochemicals as the generation of 3D structures failed for the remaining 57 phytochemicals in the database. Lastly, chemical structure of each phytochemical in SMILES, InChI and InChIKey formats was also generated using OpenBabel^63^.

Using ClassyFire (http://classyfire.wishartlab.com/)^30^, the chemical classification for each phytochemical into hierarchical levels namely, kingdom, superclass, class and subclass, was predicted. Further, using NP classifier (https://npclassifier.ucsd.edu/)^28^, a natural product specific chemical classification for each phytochemical into biosynthetic pathway, superclass and class was predicted. For each phytochemical in our database, external links to other standard chemical databases are provided using UniChem^65^. Lastly, the natural product likeness or NP-Likeness score for each phytochemical was computed using a custom RDKit script^29,31^.

For each phytochemical in our database, the physicochemical properties and drug-likeness scores were computed using in-house custom RDKit scripts. Further, the Absorption, Distribution, Metabolism, Excretion and Toxicity (ADMET) properties of the phytochemicals were predicted using SwissADME (http://www.swissadme.ch/)^66^. Since the SwissADME restricts the input molecules based on their length of SMILES, therefore, ADMET predictions could not be obtained for 493 phytochemicals in our database. Finally, we computed 1875 molecular descriptors, both 2D and 3D descriptors, for each phytochemical in our database using PaDEL^67^ software.

The predicted human target proteins of phytochemicals were obtained from the STITCH database (www.stitch.embl.de)^34^. Only high confidence phytochemical - human target protein interactions with a score of at least 700 were retrieved from the STITCH database. Further, the genes corresponding to the target human proteins were mapped to the HUGO Gene Nomenclature Committee (HGNC) symbols and identifiers^68^.

Phytochemicals with published experimental evidence of acting as covalent inhibitors were identified and compiled from CovalentInDB (http://cadd.zju.edu.cn/cidb/)^69^ and CovPDB (http://drug-discovery.vm.unifreiburg.de:8000/covpdb/)^70^ via a comparison of the chemical structures followed by manual verification.

### Molecular complexity

Molecular complexity of the phytochemicals in IMPPAT 2.0 was compared with four chemical spaces namely, phytochemicals in IMPPAT 1.0 and three collections of small molecules obtained from Clemons *et al.*^38^ corresponding to 6152 commercial compounds (CC), 5963 diversity-oriented synthesis compounds (DC’) and 2477 natural products (NP). For each compound in the above-mentioned five chemical spaces, we computed using RDKit^64^ two size-independent metrics namely, stereochemical complexity which is the fraction of stereogenic carbon atoms in a compound, and shape complexity which is the ratio of sp^3^-hybridized carbon atoms to the total number of sp^2^- and sp^3^-hybridized carbon atoms in a compound, and six other physicochemical properties namely, molecular weight, log P, topological polar surface area, number of hydrogen bond donors, number of hydrogen bond acceptors, and number of rotatable bonds.

### Molecular scaffold

Based on definition by Lipkus *et al.*^24,25^, molecular scaffolds were computed at three levels namely, graph/node/bond (G/N/B) level, graph/node (G/N) level, and graph level using RDKit^71^. Scaffolds were computed by modifying the MurckoScaffold.py from RDKit^71^. Scaffold at G/N/B level has connectivity, element and bond information, at G/N level has connectivity and element information but ignores bond information, and at graph level has only connectivity information^24,25^.

### Quantifying and visualizing chemical similarity

Chemical structure similarity between any two molecules is quantified using the widely-used metric, Tanimoto coefficient (Tc)^35^, which was computed using Extended Circular Fingerprints (ECFP4) as implemented in RDKit^71^. Chemical similarity network (CSN) consists of nodes corresponding to phytochemicals and edges connecting pairs of nodes with Tc ≥ 0.5. The value of Tc for a pair of molecules in the CSN gives the extent of chemical similarity between them, and this is captured by the thickness of the corresponding edge (Figure 10a). The maximum common substructure (MCS) for phytochemicals in a connected component of the CSN was computed using FindMCS function in RDKit^71^. The SMARTS for a MCS was visualized using SMARTSview webserver^72,73^ (https://smartsview.zbh.uni-hamburg.de/).

### Web-interface and database management

IMPPAT 2.0 database has a user-friendly web-interface and can be accessed at https://cb.imsc.res.in/imppat. The website is also mirrored at https://www.imppat.com/ and https://www.imppat.in/. The website is hosted on a local Apache (https://httpd.apache.org/) server running on Debian 9.1.3 Linux operating system. The association tables are stored in SQL format created using the open source relational database management system MariaDB (https://mariadb.org/). The front-end of the website was created using the open source CSS framework Bootstrap 4.1.3 (https://getbootstrap.com), customized with in-house HTML, PHP (http://php.net/), CSS, JavaScript and jQuery (https://jquery.com/) scripts. Further, Cytoscape.js (http://js.cytoscape.org/) is incorporated for visualizing networks, and jQuery plug-in DataTables (https://datatables.net/) for displaying tables. Also, JSME Molecule Editor^74^ is incorporated to enable drawing of chemical structures and JSmol (http://jmol.sourceforge.net/) to visualize 3D chemical structures.

### Data availability

IMPPAT 2.0 database on phytochemicals of Indian medicinal plants is accessible via the associated website: https://cb.imsc.res.in/imppat. The compiled information in IMPPAT 2.0 is made available under a Creative Commons Attribution-NonCommercial 4.0 (CC BY-NC 4.0) International License (http://creativecommons.org/licenses/by-nc/4.0/).

### Code availability

The computer codes used to analyze the phytochemical space of IMPPAT 2.0 are available via the associated GitHub repository: https://github.com/asamallab/imppat2.

## Acknowledgements

We thank B.S. Karthikeyan, Gaurav Kumar, Kishan Kumar, Geetha R and G. Rajesh for their help in data collection. We thank D. Gokul Balaji, P. Mangalapandi and B. Raveendra Reddy for computational support. Areejit Samal would like to acknowledge funding from the Department of Atomic Energy (DAE), Government of India (GoI), the Science and Engineering Research Board (SERB), GoI [Ramanujan Fellowship SB/S2/RJN-006/2014], and the Max Planck Society, Germany [Max Planck Partner Group in Mathematical Biology]. The funders have no role in study design, data collection, data analysis, manuscript preparation or decision to publish.

## Author contributions

R.P.V., K.M. and A.S. designed research. R.P.V., K.M. and A.K.S carried out the data compilation and curation. R.P.V., K.M. and A.K.S. designed the database platform and visual interface. R.P.V. performed the computational analysis. A.S. and R.P.V. wrote the manuscript. A.S. conceived and supervised the project. All authors have read and approved the manuscript.

## Supplementary Information

**Supplementary Table S1:** The table gives the list of 70 books from which Plant - Part - Phytochemical associations for Indian medicinal plants in IMPPAT 2.0 were obtained.

**Supplementary Table S2:** The table gives the list of 146 books from which Plant - Part - Therapeutic use associations for Indian medicinal plants in IMPPAT 2.0 were obtained.

**Supplementary Table S3:** The table provides the IMPPAT Phytochemical identifier, Chemical name, SMILES, InChI and QEDw score for the 1335 drug-like phytochemicals in IMPPAT 2.0 identified in this study.

